# Characterization of a flexible AAV-DTR/DT mouse model of acute epithelial lung injury

**DOI:** 10.1101/2021.06.18.445859

**Authors:** Eva Griesser, Tanja Schönberger, Birgit Stierstorfer, Hannah Wyatt, Wolfgang Rist, Thorsten Lamla, Matthew James Thomas, David Lamb, Kerstin Geillinger-Kästle

**Affiliations:** Immunology & Respiratory Diseases Research, Boehringer Ingelheim Pharma GmbH & Co. KG, Biberach an der Riss, Germany; University of Bath, Bath, United Kingdom; Drug Discovery Sciences, Boehringer Ingelheim Pharma GmbH & Co. KG, Biberach an der Riss, Germany

**Author notes:** Correspondence: Kerstin Geillinger-Kästle (Immunology & Respiratory Diseases Research, Boehringer Ingelheim Pharma GmbH & Co. KG, Biberach an der Riss, Germany). shared first author (Eva Griesser, Tanja Schönberger). **Author’s Contributions:** EG, TS, HW and KGK planned and performed experiments and analysed the data. TL provided AAV vectors. BS, WR, MJT, DL and KGK designed research. EG, TS and KGK wrote the manuscript. All authors reviewed and edited the manuscript.

**Keywords:** diphtheria toxin, DTR, AAV, epithelial injury, lung injury

## Abstract

**Background & aim:** Recurring epithelial injury and aberrant repair are considered as a major driver of idiopathic pulmonary fibrosis (IPF) leading to chronic inflammation, fibroblast activation and ultimately to scarring and stiffening of the lung. As decline of lung function is the first reported symptom by IPF patients and occurs once fibrosis is firmly established, animal models are required to study early disease-driving mechanisms.

**Methods:** We developed a novel and flexible mouse model of acute epithelial injury based on adeno-associated virus (AAV) variant 6.2 mediated expression of the human diphtheria toxin receptor (DTR). Following intratracheal administration of diphtheria toxin (DT), a cell-specific death of bronchial epithelial and alveolar epithelial type II cells can be observed.

**Results:** Detailed characterization of the AAV-DTR/DT mouse model revealed increasing cell numbers in bronchoalveolar lavage (BAL; macrophages, neutrophils, and atypical cells) and elevation of apoptotic cells and infiltrated leukocytes in lung tissue, which were dependent of viral genome load and DT dose. Cytokine levels in BAL fluid showed different patterns dependent of viral genome load with IFNγ, TNFα, and IP-10 increasing and IL-5 and IL-6 decreasing, while lung function was not affected. Additionally, laser-capture microdissection-based proteomics of bronchial and alveolar epithelium showed upregulated immune and inflammatory response in all epithelial cell regions and extracellular matrix deposition in infiltrated alveoli, while proteins involved in pulmonary surfactant synthesis, alveolar fluid clearance and alveolar-capillary barrier were downregulated in the parenchyma.

**Conclusion:** Our novel AAV-DTR/DT model resembles specific aspects of pulmonary diseases like IPF and acute respiratory distress syndrome.

**Short summary for social media:** A novel and flexible mouse model of acute epithelial lung injury based on AAV-mediated expression of the human diphtheria toxin receptor followed by intratracheal instillation of diphtheria toxin resembles specific aspects of pulmonary diseases like IPF.

## Introduction

Lung epithelial injury under certain environmental and genetic pressures leads to different pathologies that can be associated with severe outcome and increased mortality (1, 2). Chronic pathologies such as idiopathic pulmonary fibrosis (IPF) or more acute pathologies such as acute respiratory distress syndrome (ARDS) or IPF exacerbation both exhibit a breakdown of the epithelial barrier and impairment of lung function (1, 3, 4). Repeated epithelial injury can lead to chronic inflammation, fibroblast activation and ultimately to scarring and stiffening of the lung. Particularly in the chronic setting, the observed decline in lung function is often the first symptom reported by IPF patients. As these changes occur slowly over time, patients only present to clinicians at a progressed state of the disease. Therefore, disease driving mechanisms can only be estimated based on observed changes in biomarkers, the transcriptome or proteome. These changes indicate a strong epithelial impact on disease progression as epithelial biomarkers like pulmonary surfactant proteins D and A (SP-D, SP-A) as well as KL-6 are found to be increased in plasma as well as associated with disease prognosis (1, 4, 5). In addition, single cell sequencing showed marked changes in epithelial cell type composition and gene expression (6).

Currently being unable to identify patients early in the disease phase, alterations in epithelial cell signalling pathways at a very early stage of disease development can only be investigated *in vitro* or in animal models. In literature, several models of lung injury are described: the genetically modified DTR/DT model (diphtheria toxin receptor/diphtheria toxin), ventilator induced lung injury, smoke exposure, lipopolysaccharide (LPS) administration, bleomycin, or infectious models, each of which address specific aspects of lung injury.

While smoke exposure, ventilator induced lung injury, LPS as well as infectious models use the main disease relevant stimuli, models such as the bleomycin induced lung fibrosis mainly resemble the mechanistic aspects of the human disease. This is also the case for the widely used DTR/DT cell depletion system. The human DTR (hDTR) is sensitive for diphtheria toxin (DT), thereby inducing apoptosis by blocking protein synthesis. In contrast, a polymorphism of the murine DTR protects mice from DT mediated toxicity (7). Since the first description of genetic DTR mice a variety of genetic mouse strains have been generated expressing the human DTR in specific cell types such as neutrophils or regulatory T-cells (8, 9) and used in various research fields.

However, the generation of cell type or organ specific DTR mice is costly and time consuming. Therefore, our aim was to develop a novel and flexible DTR/DT model based on adeno-associated virus (AAV) vector mediated hDTR expression. We comprehensively characterized the AAV6.2 driven lung specific cell depletion model 24 h after dose dependent injury. Dose dependency was achieved by increasing the viral genome (vg) load or administered DT dose. This allowed equal amount of cell depletion, by increasing hDTR expression while reducing DT instillation and thereby minimising hDTR independent effects of DT. Observed changes in cytokine secretion, immune influx and protein regulation resemble hallmarks of IPF. In addition, region specific responses to injury were detected using laser-capture microdissection-based proteomics.

## Methods

Detailed methods can be found in the online supplemental data.

### Adeno-associated virus (AAV) to overexpress hDTR

The expression cassette consisting of a Kozak sequence followed by the coding sequence for human diphtheria toxin receptor (hDTR; UniProt Q53H93) was codon-usage optimized for expression in mice and synthesized (Thermo Fisher Scientific/GeneArt, Regensburg, Germany) before being cloned into a pFB vector. The vector contained AAV2 inverted terminal repeats, from which one was lacking the terminal resolution site, a cytomegalovirus (CMV) promoter and a SV40 polyA signal. As a control a pFB-stuffer vector (contains non-coding “ stuffer” DNA) (29) was used. Recombinant AAV6.2 vectors were produced by calcium phosphate transfection of human embryonic kidney (HEK)-293 cells using the pFB_CMV-hDTR or the pFB_stuffer plasmid in combination with a pAAV-Cap6.2 (56) and the pHelper plasmid (Thermo Fisher Scientific), followed by purification and titer determination as described previously (57).

### Animals

Male C57BL/6JRj mice at the age of 8-12 weeks were purchased from Janvier Labs (France). Mice were housed in groups of 2-5 mice in individually ventilated cages at 22-25 °C, at a humidity of 45-65%, and at a 12 h day/night cycle with free access to water and food. Ethical approval was obtained from the regional board for animal care and welfare (Regierungspräsidium Tübingen, Germany, TVV-18-23-G). At day zero AAV-stuffer or AAV-hDTR was instilled intratracheally under isoflurane anaesthesia (Isofluran CP; CP-Pharma, Burgdorf, Germany). After 28 days 100 ng, 150 ng or 200 ng diphtheria toxin (DT) from Sigma-Aldrich was instilled intratracheally in all experimental groups. Animals were euthanised 24 h after DT instillation. Plasma, bronchoalveolar lavage (BAL) and lung tissue were obtained and prepared according to needs of further analysis.

### Bronchoalveolar lavage

BAL was generated by flushing the lungs twice with HANKS salt solution. After analysis of total cell influx as well as cell types using the Sysmex XT-1800i device, BAL cells were pelleted and stored at -80 °C. The supernatant (BAL fluid - BALF) was aliquoted and used for cytokine analysis.

### ELISA

All ELISA assays were performed according to manufacturer’s recommendations. SP-D was measured in plasma using the Mouse SP-D Immunoassay (#MSFPD0) from R&D systems (Abingdon, United Kingdom). Cytokine levels in BALF were detected using U-PLEX Biomarker Group 1 (ms) 35-Plex from Meso Scale Discovery (MSD; Rockville, MD).

### RT-PCR

BAL cells were lysed in RLT Plus buffer (Qiagen) and RNA was extracted using the MagMAX™ Express Magnetic Particle Processor (Thermo Scientific). 400 ng RNA was transcribed into cDNA using the High-Capacity cDNA Reverse Transcription Kit (Thermo Scientific). RNA of Nos2 (Mm00440502_m1), Tnf (Mm00443258_m1), Il6 (Mm00446190_m1), Fizz1 (Mm00445109_m1), Cxcl1 (Mm04207460_m1), and Arg1 (Mm00475988_m1) was quantified with the commercial TaqMan® Fast Advanced assays (Thermo Scientific) using a ViiA7 PCR device (Applied Biosystems).

### Histology and immunohistochemistry

After flushing the lungs for BAL, the left lobes of the lungs were pressure-filled with low melting agarose (0.75%, Sigma Aldrich) and transferred to 10% neutral buffered formalin (Sigma-Aldrich, # HT501128). Tissue was fixed for at least 24 h before samples were processed with an automated tissue processor (Tissue-Tek® VIP® 6, Sakura), embedded in paraffin and cut into 3 µm sections followed by haematoxylin & eosin, TUNEL and immunohistochemical staining as described in detail in the online supplemental methods.

## Results

### Characterization of AAV-hDTR dose dependent cell depletion *in vivo*

Four different doses of AAV-hDTR were administered intratracheally. After 28 days of expression, three different doses of DT were applied intratracheally to initiate cell depletion. 24 h after administration, the effects of DT were characterized. With increasing viral genome (vg) load and consequently higher hDTR expression, 100 ng DT led to increased epithelial damage, as measured by rising levels of pulmonary surfactant protein D (SP-D) (Fig. 1A). To confirm the expression of hDTR in the murine lung after virus treatment, *in situ* hybridization (ISH) was performed investigating expression of hDTR-mRNA in bronchial and alveolar epithelial cells (Fig. S1A). In addition, double staining for hDTR-mRNA (ISH) and apoptotic cells (TUNEL) showed colocalization in affected cells (Fig. S1B/D). In contrast, in an AAV-stuffer control treated lung, administered with an AAV vector containing non-coding “ stuffer” DNA (10), only few apoptotic cells could be seen (Fig. S1C). However, epithelial cell death did not affect the lung function in the AAV-DTR/DT model (Fig. S2). Using a flow cytometric haematology system, an increase in neutrophils and macrophages was detected with higher viral genome load and with increasing DT doses in bronchoalveolar lavage (BAL) (Fig. 1C/D). In addition, elevated numbers of atypical cells were detected (Fig. 1B). These cells showed lower granularity and reduced size detected with side scatter and forward scatter in the flow cytometric haematology analysis, suggesting that these cells undergo cell death. Immune cell infiltration as well as epithelial cell death were confirmed by (immuno-) histology. Morphological evaluation of H&E-stained sections showed degenerating cells characterized by pyknotic nuclei and apoptotic bodies (Fig. S3) while epithelial coverage remained yet intact. TUNEL staining revealed the extent of apoptosis in bronchial and alveolar epithelium dependent on the viral genome load (Fig. 2A/C; Fig. S4). Perivascular and peribronchiolar cell infiltration seen in H&E-stained sections was confirmed by CD45 immunohistochemistry (IHC) staining showing leukocyte infiltration in a virus load dependent manner (Fig. 2B/C; Fig. S4).

**Figure 1:**
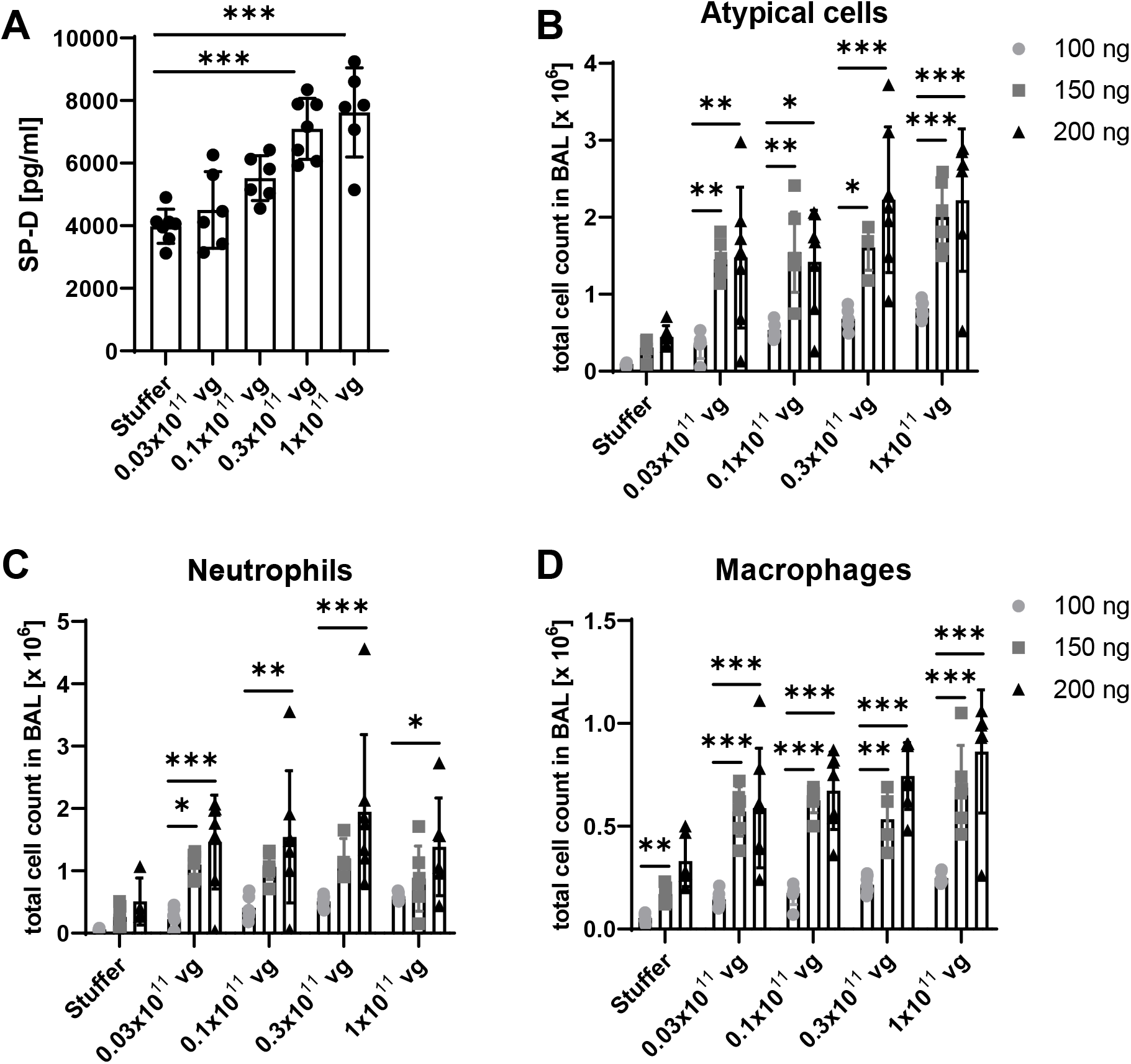
Intratracheal DT application induces epithelial cell death and immune infiltration. (A) SP-D leakage into plasma of AAV-stuffer (1 ⨯ 10^11^ vg) and AAV-hDTR treated animals 24 h after administration of 100 ng DT. Number of (B) atypical cells, (C) neutrophils and (D) macrophages detected in BAL 24 h after administration of indicated DT doses using the Sysmex flow cytometric haematology analyser. Two-way ANOVA followed by Tukey’s multiple comparisons test was applied (n=5-7, *p <0.05, **p <0.01, ***p <0.001).

**Figure 2:**
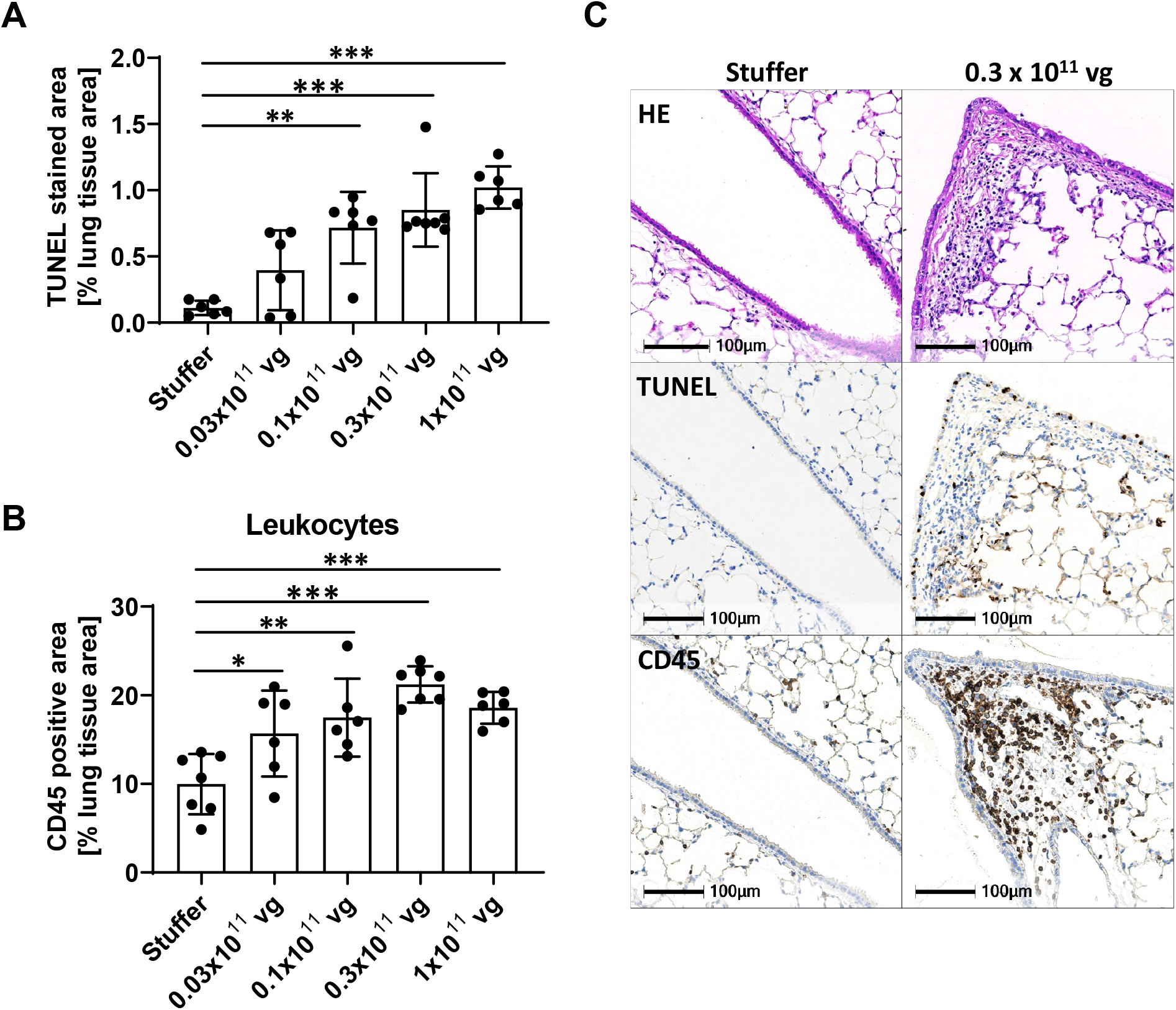
Intratracheal DT application induces epithelial cell death and immune infiltration. (A) Quantitative analysis of TUNEL stained lung sections (apoptosis) from AAV-stuffer (1 × 10^11^ vg) and AAV-hDTR treated animals which received 100 ng DT. (B) Quantification of CD45-staining (leukocytes) confirms increasing immune infiltration with higher viral genome (vg) load. One-way ANOVA with Dunnett’s multiple comparisons test was applied (n=5-7, *p <0.05, **p <0.01, ***p <0.001). (C) Representative images of H&E-stained lung sections (upper panel) show morphological signs of epithelial cell death in AAV-hDTR treated lungs (0.3 ⨯ 10^11^ vg), which was confirmed by TUNEL staining (middle panel). Subsequent CD45-stained sections (lower panel) show peribronchiolar infiltrating leukocytes in the affected area. Bronchial and alveolar airways contain only occasional inflammatory cells as bronchoalveolar lavage and agarose instillation of lungs were performed before further tissue processing.

DTR/DT mediated cell depletion relies on the expression of the human DTR (HBEGF), as the murine DTR (Hbegf) has no affinity to DT. Upon binding of DT to its receptor, it is internalized via endocytosis and split into fragment A (DT-A) and fragment B (DT-B). Once DT-A is released into the cytosol it blocks protein synthesis by inhibiting polypeptide chain elongation factor 2, finally leading to cell death (7). However, we observed also in AAV-stuffer treated mice, which do not express the human DTR, a DT dose dependent effect. An increase in macrophages and in neutrophils was detected in the BAL (Fig. 3A). Elevated number of immune cells resulted also in higher concentrations of macrophage inflammatory proteins (MIPs) (Fig. 3B). In addition, after intratracheal administration of 150 ng and 200 ng DT atypical cells were detected in the BAL (Fig. 3A). However, observed increase in atypical cells did not result in enhanced leakage of SP-D from the epithelial lining fluid into the plasma (Fig. S5), suggesting unspecific effects of DT at higher doses. These unspecific effects of DT in mice limit available genetic DTR/DT models.

**Figure 3:**
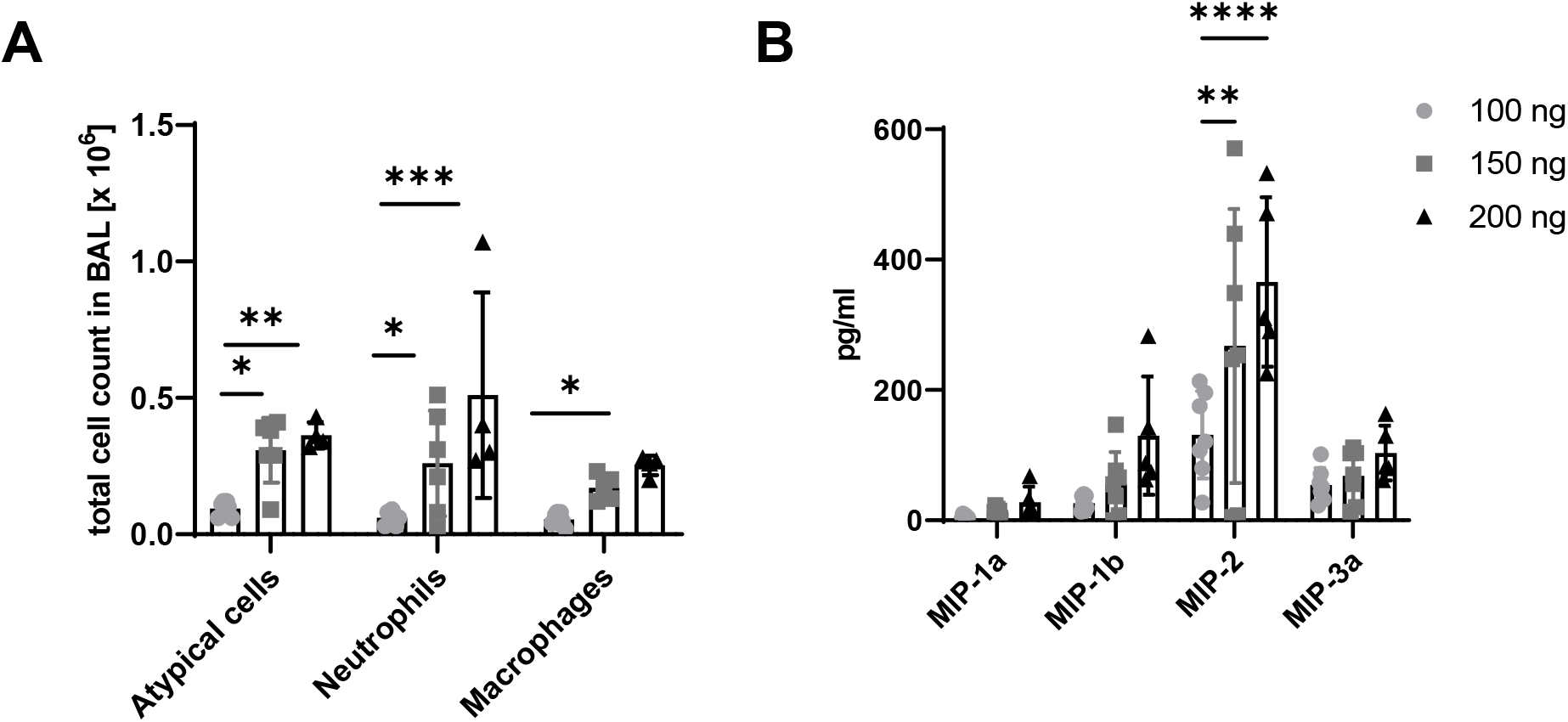
Intratracheal DT application induces immune infiltration and cytokine release independent of hDTR expression. (A) Total cell counts in BAL and (B) cytokine release in BALF 24 h after DT administration of indicated doses in AAV-stuffer treated mice (1 ⨯ 10^11^ vg). Two-way ANOVA followed by Tukey’s multiple comparisons test was applied (n=5-7, *p <0.05, **p <0.01, ***p <0.001, ****p<0.0001).

Our data implies that increasing the expression of hDTR can compensate for lower DT application to limit unspecific effects. Consequently, 100 ng DT was used for subsequent studies.

### Characterization of cytokines in BALF

Cytokine concentrations were evaluated using a 35-plex MSD panel for all four AAV-hDTR dosages (Table S1). Interestingly, cytokine profiles showed three different patterns with increasing viral genome load (Fig. 4). GM-CSF and KC/GRO strongly decreased upon cell death, indicating the importance of epithelial cells for their production and secretion (Fig. 4A). Most detectable cytokine concentrations elevated with increasing injury (e.g. IL-12, IL-16, and TNFα; Fig. 4B and Table S1). However, some cytokines showed increased levels at low viral genome load but decreased with higher viral load (e.g. IL-5, IL-6, and VGEF; Fig. 4C). In addition, we observed increased concentrations of IFNγ and IP-10 as well as MIP-1b and MIP-3a (Fig. 5). Correlation matrix analysis resulted as expected in a strong correlation of IP-10 and IFNγ (Pearson correlation coefficient 0.9; Fig. S6). Furthermore, TNFα levels correlated with IL-12, MIP-1b (both coefficient 0.9) and MCP-1 (coefficient 0.7). Based on these data, we subsequently analysed macrophage polarization markers in BAL cells.

**Figure 4:**
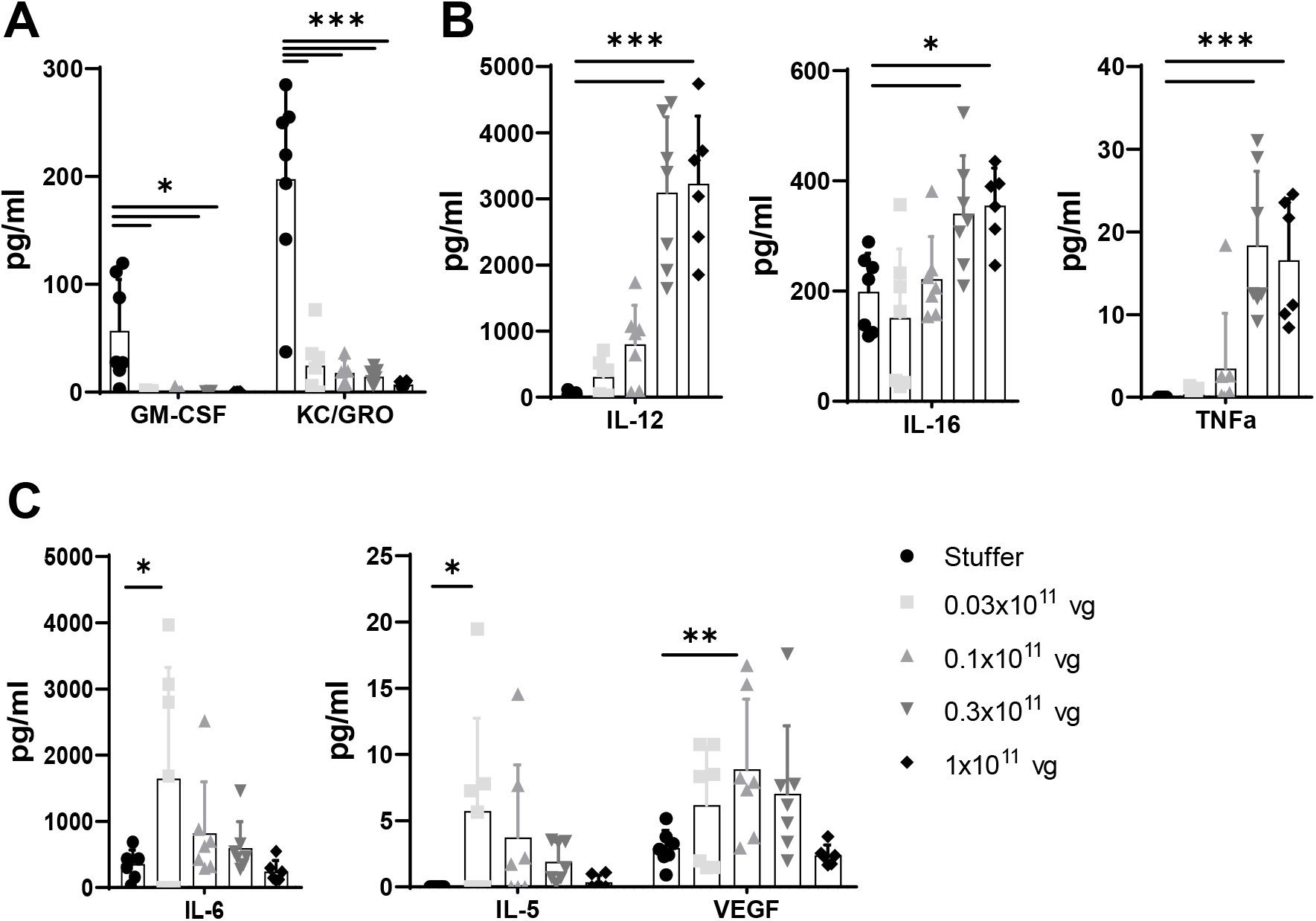
Cytokines in the BALF of AAV-stuffer (1 ⨯ 10^11^ vg) and AAV-hDTR treated animals upon epithelial cell death induced by intratracheal administration of 100 ng DT. (A) GM-CSF and KC/GRO decreased upon acute epithelial lung injury. (B) IL-12, IL-16 and TNFα increased, while (C) IL-5, IL-6 and VEGF decreased with higher viral genome (vg) load. One-way ANOVA followed by Dunnet’s multiple comparisons test was applied (n=5-7, *p <0.05, **p <0.01, ***p <0.001).

**Figure 5:**
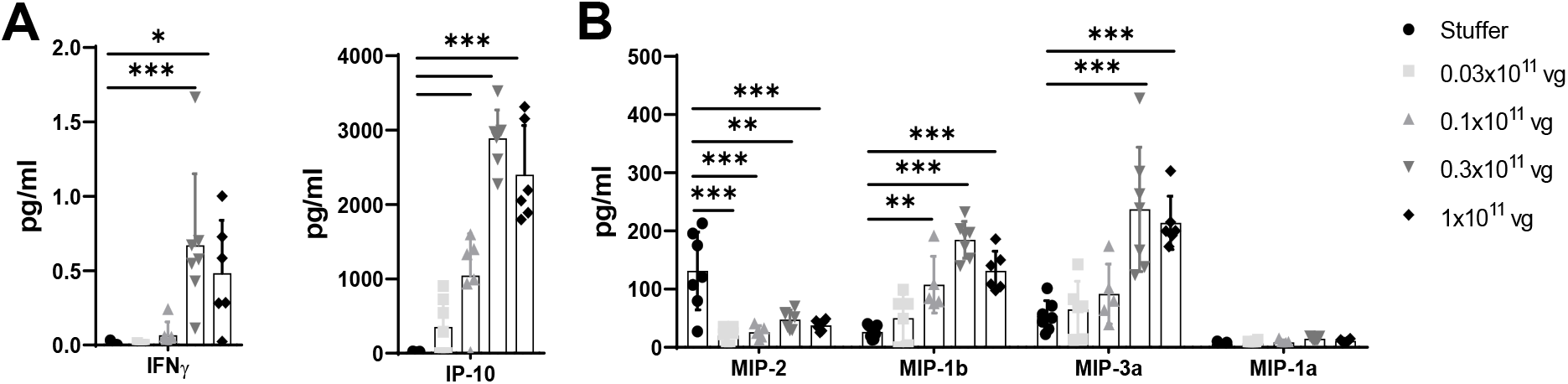
Cytokines in the BALF of AAV-stuffer (1 ⨯ 10^11^ vg) and AAV-hDTR treated animals upon epithelial cell death induced by intratracheal administration of 100 ng DT. (A) IFNγ and IP-10 as well as (B) MIP-1b and MIP-3a increased, while MIP-2 decreased with higher viral genome (vg) load. Two-way ANOVA followed by Tukey’s multiple comparisons test was applied (n=5-7, *p <0.05, **p <0.01, ***p <0.001).

### M1 and M2 macrophage markers increase dependent on viral genome load

Upon activation by various stimuli macrophages can be polarized into different subtypes. Typically, the first hierarchical division is into M1 macrophages being considered as inflammatory macrophages and M2 macrophages presenting anti-inflammatory/pro-repair characteristics (11, 12). Remarkably, we identified markers for both macrophage subtypes, which showed increase at the highest AAV-hDTR doses. However, M1 markers iNos, Tnf, and Il-6 (Fig. 6A) were already upregulated with the second highest viral genome load (0.3 ⨯ 10^11^ vg) in contrast to M2 markers Fizz1 and Cxcl1, which were only significantly elevated with 1 ⨯ 10^11^ vg, while Arg1 levels didn’t change (Fig. 6B).

**Figure 6:**
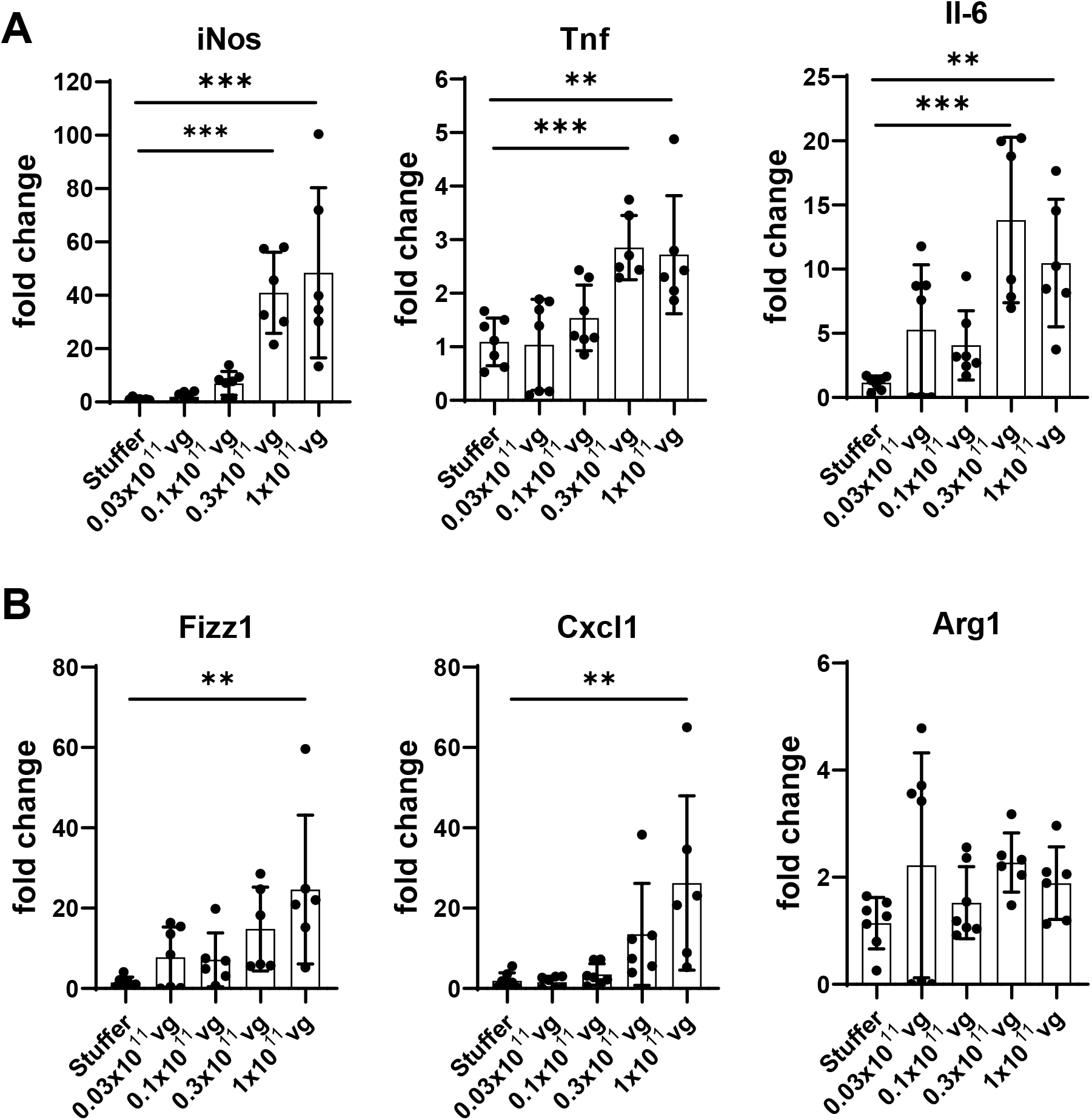
mRNA levels of macrophage polarization markers detected in BAL cells of AAV-stuffer (1 ⨯ 10^11^ vg) and AAV-hDTR treated animals administered with 100 ng DT. mRNA of (A) M1 polarization markers iNos, Tnf and Il-6 and (B) M2 polarization markers Fizz1, Cxcl1 and Arg1 was measured. One-way ANOVA followed by Dunnet’s multiple comparisons test was applied (n=5-7, *p <0.05, **p <0.01, ***p <0.001).

### Evaluation of the epithelial proteome using laser-capture microdissection

Different regions of formalin-fixed and paraffin-embedded (FFPE) lung tissue were isolated using laser-capture microdissection to analyse proteomic changes in the bronchial and alveolar epithelium (Fig. S7). The analysis of total three batches with 14 samples per batch included tandem mass tagging and high-resolution mass spectrometry as described recently (13). Samples from the same lung region (bronchi, alveoli) were analysed in the same batch for direct comparison of AAV-hDTR (0.3 ⨯ 10^11^ vg) to AAV-stuffer mice. In the third batch immune cell infiltrated and normal alveolar tissue from AAV-hDTR mice were combined.

Analysis of the bronchial epithelium presented a clear difference between hDTR and stuffer mice (Fig. S8A). 46 proteins were significantly upregulated (fold change ≥1.5, BH-adj. p-value <0.05; Fig. S8B, tables S2/S5), which are mainly involved in immune response, cellular response to interferon-beta and defence response to virus (Fig. S8C). Interestingly, only five proteins were downregulated in the bronchial epithelium of AAV-hDTR mice upon DT instillation including secretory proteins Scgb1a1, a marker for club cells, Hp and Wfdc2 as well as Dpep1 and Kndc1 (Fig. S8B).

Combined data analysis of both alveolar tissue batches resulted in a clear differentiation between normal tissue in stuffer and hDTR mice and infiltrated tissue in hDTR mice (Fig. 7A). Hierarchical clustering of proteins with a significant change in their quantities between at least two experimental groups (one way ANOVA BH-adj. p-value <0.05) resulted in clusters of proteins with increased protein quantities in the infiltrated tissue compared to the normal one (cluster 4-6; Fig. 7B). Among these proteins, innate immune response, complement activation, translation, transcription, and mRNA processing were overrepresented (Fig. 7C). Cluster 1-3 include proteins with lower protein quantities in the alveolar tissue from hDTR mice. Extracellular matrix organization, cell adhesion as well as mitochondrial metabolic processes such as TCA cycle and fatty acid metabolism were overrepresented in these clusters. Alveolar epithelial type II (ATII) cell markers SP-C, SP-A and Muc1 were significantly downregulated during acute epithelial injury confirming depletion of ATII cells (Fig. 7D, tables S3/S6). Additionally, proteins from the fatty acid and lipid metabolism, such as Lpcat1, Fabp5 and Acsl4, and proteins being part of the lung fluid transport, Aqp5 and Atp1b1, were significantly downregulated in normal alveolar tissue isolated from hDTR mice compared to stuffer mice upon DT instillation. On the other hand, extracellular matrix proteins, including all three fibrinogen subunits (Fga, Fgb, Fgg) and two integrin subunits (Itgam, Itgb2), presented increased levels. Comparison of infiltrated with normal tissue from hDTR mice also resulted in significant upregulation of proteins from the interstitial extracellular matrix including five collagen proteins (Col1a1, Col1a2, Col3a1, Col5a1, Col5a2), Eln and Bgn (Fig. 7D, tables S4/S7). Conversely, proteins from the basement membrane, e.g. Col4a3 and Agr, decreased in infiltrated tissue (Fig. 7D).

**Figure 7:**
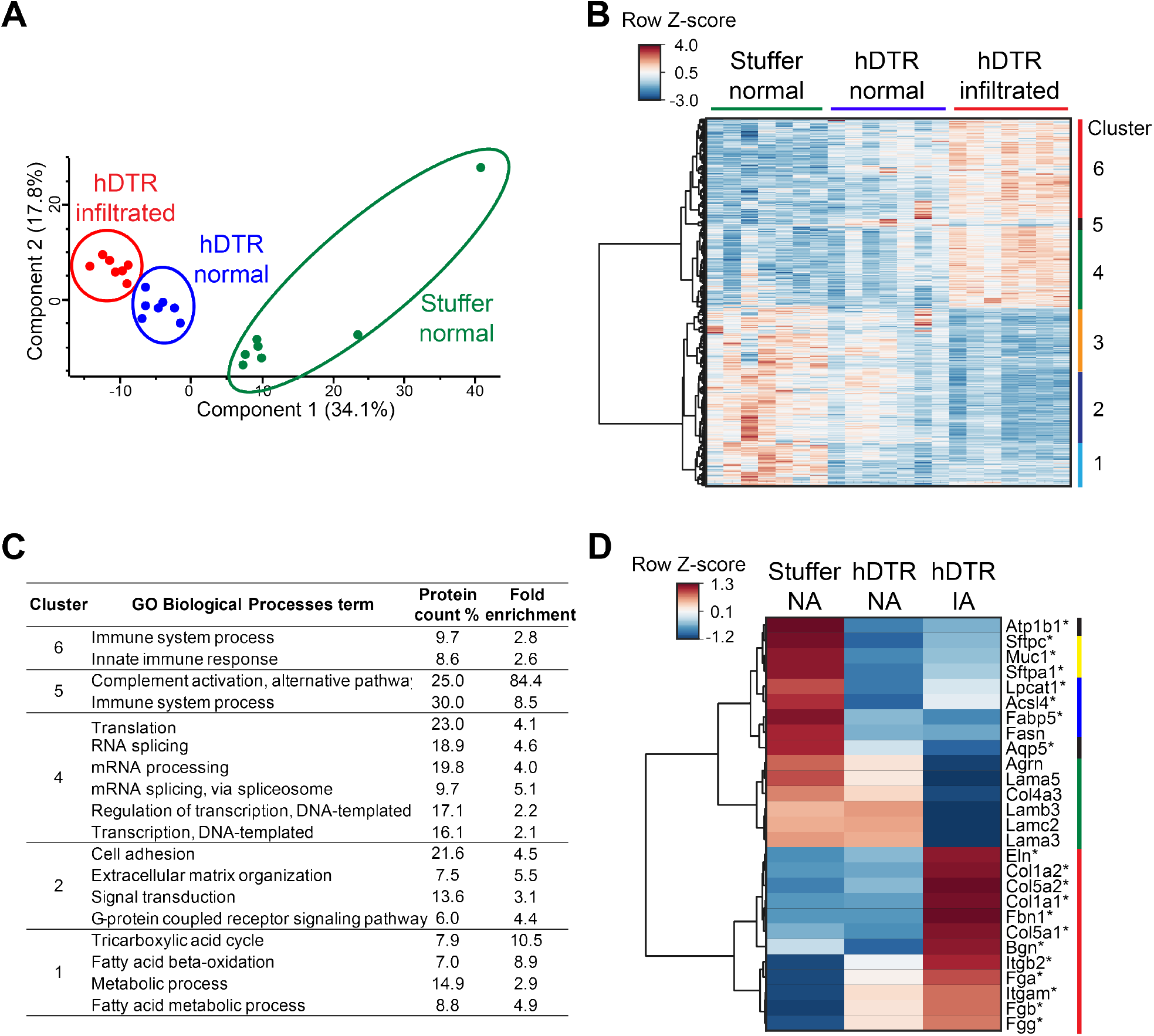
Spatial resolution of proteomic changes in alveolar tissue upon acute epithelial injury. (A) Principal component analysis scores plot of log2-transformed protein quantities of normal alveolar tissue from AAV-hDTR (0.3 ⨯ 10^11^ vg) and AAV-stuffer (1 ⨯ 10^11^ vg) mice and infiltrated tissue from AAV-hDTR mice 24 h after intratracheal DT application (100 ng). (B) Hierarchical clustering depicting row Z-scores of protein quantities from three experimental groups (n=7; only proteins with one-way ANOVA Benjamini Hochberg (BH) adjusted p-value <0.05 were considered). (C) Enriched Gene Ontology (GO) Biological Processes annotated to proteins from each of six clusters shown in panel B. Alveoli-specific proteome was used as background (Fisher’s exact test, BH-corrected p-value <0.05, protein count ≥6%). (D) Heatmap showing the average row Z-scores (n=7) of selected proteins involved in interstitial extracellular matrix (red), basement membrane (green), fatty acid/lipid metabolism (blue) and lung fluid transport (black) as well as ATII cell markers (yellow). *Proteins were significantly deregulated (fold change ≥1.5, BH-adjusted p-value <0.05) when comparing hDTR normal to stuffer normal or hDTR infiltrated to hDTR normal. NA – normal alveolar tissue, IA – infiltrated alveolar tissue. Protein quantities are the summed TMTpro reporter ion signal-to-noise ratios of corresponding peptides.

## Discussion

In this study we developed a novel and flexible DTR/DT mouse model based on AAV-mediated expression of the human DTR. Specifically, the utilized AAV variant 6.2 transduces into bronchial and alveolar epithelial type II cells. Intratracheal DT application therefore induces acute cell depletion of these cell types mimicking epithelial lung injury. In-depth characterization of the new model demonstrated increase of immune cell influx, apoptosis, SP-D leakage into plasma, and higher cytokine levels in the BALF like IFNγ, TNFα, and IP-10 with higher viral genome load. Additionally, decrease of cytokines like GM-CSF, KC/GRO, and MIP-2 was shown. Also, spatially-resolved analysis of the proteome of different epithelial cell regions revealed the impact of immune cells on extracellular matrix deposition upon injury.

### DT off-target effects

The DTR/DT system is widely used to deplete various cell types to study their role for example in immune response, repair, and regeneration of disease relevant animal models (14–16). Several genetically modified animals are available expressing the human DTR in different cell types and tissues (www.informatics.jax.org). However, there is very limited knowledge on the hDTR independent effects of DT *in vivo* in general, although there are increasing numbers of publications using the DTR/DT cell depletion system in combination with systemic (e.g. intraperitoneal) or local (e.g. subcutaneous, intratracheal) administration (17–20). Using three different DT doses in AAV-stuffer control animals we measured a DT dose dependent influx of neutrophils and macrophages and correspondingly also increased concentrations of MIPs. In addition, intratracheal DT application in AAV-stuffer treated mice seemed to also induce apoptosis in a small fraction of cells independent of hDTR expression. Similar effects were observed even 19 days after intratracheal administration of 50 ng of DT (17). However, these animals underwent OVA sensitization and challenge. Another group investigated the lung function after intratracheal application of 100 ng DT and observed changes in total lung resistance and elastance 96 h after DT administration (21). These observations were not confirmed in our experiments 24 h after DT administration and might therefore be detectable only at later timepoints. In conclusion, the hDTR independent effects of DT need to be considered when developing new models utilising specific routes of DT administration. For example, increasing hDTR expression might be a possibility to decrease the DT dose and thereby limit hDTR independent effects.

### Immune cell response to lung epithelial injury

Nonetheless, we observed epithelial cell death specifically in AAV-hDTR treated mice. Epithelial layers, which are in regular contact with environmental insults such as pathogens and environmental particles, have developed a complex response of resident epithelial and immune cells as well as infiltrating immune cells to the insult (22). Thereby resident epithelial and immune cells secrete chemokines and cytokines to attract immune cells to the site of injury. As expected upon the sterile injury leukocytes including neutrophils and macrophages were attracted by released chemokines and cytokines, such as TNFα and IFNγ. However, chemokines such as KC/GRO and MIP-2, known chemoattractants for neutrophils (23), were decreased 24 h post-injury, indicating that KC/GRO and MIP-2 are not the major drivers of immune cell influx upon epithelial injury. Induced cell death might even be the main cause for the decline as both chemokines are also secreted by lung epithelial cells (23). However, the decrease in MIP-2 might also be due to a T helper (Th) 1 cell dominance in the AAV-DTR/DT model. Mueller et al. showed that IFNγ polarizes naïve T-cells towards Th1 cells, while suppressing Th2 cell polarization (24) resulting in decreased MIP-2 and elevated MIP-1 and MIP-3 levels. This specific pattern of MIPs was also observed in the BALF of AAV-DTR/DT mice. In contrast, Th2 specific IL-4 and IL-13 (25) were not detected. Other potent stimulator of Th1 cell differentiation besides IFNγ are IL-12 and IP-10, which were all increased in the BALF of AAV-DTR/DT mice. Remarkably, elevated levels of IL-12 have also been shown in IPF patients (26), whereas upregulation of IFNγ was demonstrated specifically in T cells from IPF lungs by single cell RNAseq analysis (27). Interestingly, Dixon et al administered activated Th1 cells intratracheally to naïve mice, which resulted in increased IP-10 levels amplifying the Th1 inflammatory cell response in a positive feedback loop (28). While compiled data indicates a clear polarization of Th cells, this was not observed for macrophages, when BAL cells were analysed for M1 and M2 markers. Although M1 markers seemed to increase already at lower insults, M2 marker (except for Arg1) were also significantly upregulated.

LCM-proteomics analysis showed increase in immune and inflammatory response in both bronchial and alveolar epithelial cell regions. Specifically, a strong interferon mediated immune signature (upregulation of e.g. Stat1, Mnda, Ifit1, Ifit3) could be detected, which was consistent with BALF cytokine measurements. This response is solely induced by cell depletion of epithelial cells and not by AAV6.2 vectors, as their low immunogenicity was demonstrated previously (10, 29). Furthermore, there are no pathogen-associated stimuli involved in this model, which could trigger the observed immune reaction. In the infiltrated parenchyma Vcam1 was upregulated, which has been shown to be fundamental for neutrophil as well as monocyte migration into the lung (30, 31) and was also found to be elevated in lung tissue as well as plasma of ILD patients (32).

### LCM-proteomics reveals impact of immune cells on deposited matrix composition

Besides upregulation of Vcam1 increased levels of proteins from the interstitial extracellular matrix, such as collagens type I, III and V, elastin and biglycan were observed in immune cell infiltrated alveolar tissue. Extracellular matrix synthesis and deposition is an early event after epithelial injury and immune cell infiltration and part of the repair and wound healing process by providing a scaffold for cell migration to the site of injury (33). As leukocyte infiltration of the tissue occurs perivascular, infiltrated alveolar tissue was microdissected in perivascular proximity in a way that a small part of the infiltrate was isolated together with the neighbouring alveolar tissue (see Fig. S7). Thus, samples contained (besides alveolar cells) small portions of immune cells and extracellular matrix surrounding blood vessels. As infiltrated tissue was compared to normal tissue, which was not isolated from perivascular regions, we cannot rule out that an observed increase in levels of extracellular matrix proteins is related to these differences in isolated areas. However, with all three fibrinogen subunits (α, β, γ) extracellular matrix proteins were already elevated in normal alveoli from the AAV-DTR/DT mouse model. Fibrinogen is part of the provisional matrix, which is formed following injury to facilitate cell migration (34). Adhesion of isolated ATII cells to fibrinogen (35) and secretion of fibrinogen from A549 cells treated with IL-6 and dexamethasone were shown previously (36). Additionally, two integrin subunits (αM, β2) were increased in normal alveolar tissue, which form the leukocyte integrin αMβ2 and promote cell migration via cell adhesion to fibrinogen (37).

Basement membrane proteins decreased in the infiltrated region including collagen type IV, laminin subunits and agrin. They are part of the alveolar-capillary barrier, which consists of alveolar epithelium, capillary endothelium, and the alveolar basement membrane (38). Damage to the alveolar-capillary barrier leads to increased permeability and is a hallmark of acute lung injury and ARDS (39, 40). Clearance of the alveolar lining fluid is another critical process in the alveolar epithelium, which was affected during acute epithelial injury as shown by the downregulation of water channel Aqp5 and ion channel Atp1b1, both required for lung fluid transport (41).

The AAV6.2 variant transduces specifically into bronchial epithelial and ATII cells (10, 29). Indeed, intracellular decrease of known epithelial cell markers in the alveolar epithelium (ATII: SP-C, SP-A, Muc1) as well as bronchial epithelium (club cells: Scgb1a1) (42, 43) confirmed their depletion after DT administration. Additionally, Hp (club cells) and Kndc1 (ciliated cells), recently identified as putative epithelial cell markers by single-cell RNA sequencing (42), showed intracellular decrease in the bronchial epithelium. Besides the pulmonary surfactant proteins SP-A and SP-C proteins from the lipid and fatty acid metabolism (Lpcat1, Acsl4, Fabp5, Fasn) were decreased during acute epithelial injury, which altogether might affect the synthesis of pulmonary surfactant (44). Lpcat1 is required for the regulation of the synthesis of saturated phosphatidylcholine, the most abundant pulmonary surfactant phospholipid (45). Downregulation of lipid metabolism in ATII cells during bleomycin-induced lung injury in young and old mice at day 4 as well as in ATII cells from IPF lungs has been described recently (46).

Overall, LCM-proteomics results demonstrate matrix deposition promoting cell migration as well as decrease in alveolar-capillary barrier, alveolar fluid clearance and pulmonary surfactant synthesis in lower airways during acute epithelial injury.

### AAV-DTR/DT injury model resembles aspects of acute and chronic respiratory diseases

Characterization of the AAV-DTR/DT model revealed hallmarks of acute pulmonary diseases like ARDS as well as chronic diseases like IPF. Interestingly, there is also a great overlap of clinical biomarker of both diseases (1, 47). ARDS and IPF patients show increased SP-D levels in plasma, which is a biomarker for epithelial integrity (48, 49), underpinning the pivotal role of an intact epithelial layer. After insult to the epithelial layer ECM deposition is induced in IPF and ARDS (50, 51). In addition, both diseases show immune cell infiltration comprised of neutrophils and leukocytes as seen in the AAV-DTR/DT model (52, 53). Despite some overlap in features resembling acute and chronic lung diseases there are also specific aspects to the one or the other, like the involvement of IFNγ. While it is potentially beneficial for IPF patients it seems to be a major driver of alveolar injury in ARDS (54, 55).

Therefore, the newly developed AAV-DTR/DT model can be used to study activation of repair mechanisms in the absence of pathogen or chemical induced immune responses to better understand the onset of pulmonary diseases like ARDS and IPF.

## Abbreviations

AAV: adeno-associated virus
DTR: diphtheria toxin receptor
DT: diphtheria toxin
IPF: idiopathic pulmonary fibrosis
ARDS: acute respiratory distress syndrome
BAL: bronchoalveolar lavage
SP-A/C/D: pulmonary surfactant protein A/C/D
LPS: lipopolysaccharide
hDTR: human diphtheria toxin receptor
vg: viral genome
ISH: *in situ* hybridization
IHC: immunohistochemistry
MIP: macrophage inflammatory protein
FFPE: formalin-fixed and paraffin-embedded
ATII: alveolar epithelial type II cells

## Acknowledgements

We would like to thank Helene Lichius and Sylvia Blum for their contribution to the *in vivo* experiment as well as Anita Schoenleber, Annika Maier, Michael Schilling, and Martina Keck for excellent technical assistance.

## Data Availability

The mass spectrometry proteomics data have been deposited to the ProteomeXchange Consortium (58) via the PRIDE (59) partner repository with the dataset identifier PXD025712.

