## Supplemental data for "Characterization of a flexible AAV-DTR/DT mouse model of acute epithelial lung injury"

**shared first author*

[#]*Correspondence: Kerstin Geillinger-Kästle (Immunology & Respiratory Diseases Research, Boehringer Ingelheim Pharma GmbH & Co. KG, Biberach an der Riss, Germany, Tel.: +49 7351 54-175166,)*

### Supplemental methods

#### Lung function

Animals were anesthetized by intraperitoneal injection of pentobarbital (Narcoren®, Boehringer Ingelheim; 60 mg/kg) and xylazine (Rompun®, Bayer Vital GmbH; 2.5 mg/kg). For ventilation a cannula was inserted into the trachea and coupled to a flexiVent® small animal ventilator (SCIREQ, Montreal, Quebec, Canada) and controlled using flexiWare 7.5. Mechanical ventilation was started with a tidal volume of 6.5 mL/kg, a frequency of 150 breaths/min and a positive end-expiratory pressure of 3 $cmH_2O$.

#### Histology and immunohistochemistry

Haematoxylin and eosin (H&E) staining was performed according to standard protocols using the Leica ST5020 Multistainer (Leica Biosystems Nussloch GmbH, Nussloch, Germany). Immunohistochemical (IHC) staining and TUNEL assay were carried out on the automated Leica Bond™ platform (Leica Biosystems, Melbourne, Australia). Leukocytes were detected using an anti-CD45 antibody (#ab10558, abcam) after heat induced antigen retrieval in a concentration of 2.8 µg/µL. Apoptotic cells were detected using the TUNEL assay (DIG-11-dUTP, #11570013910, Roche; TdT-Enzyme, #M1875, Promega). Bound antibodies were visualized using the Bond™ Polymer Refine Detection System (Leica Biosystems, Newcastle, United Kingdom). Following scanning with the Axio Scan.Z1 (200x magnification; Carl Zeiss Microscopy GmbH, Jena, Germany), the CD45- and TUNEL-positive stained areas were determined using HALO™ (Indica Labs, Corrales, NM). Quantitative data were expressed as percentage of stained area relative to whole-sectional area.

#### hDTR *in situ* hybridization and TUNEL IHC assay

To confirm expression of hDTR (proheparin-binding EGF-like growth factor, HBEGF) in pulmonary cells in relation to apoptotic cells after DT administration, an *in situ* hybridization (ISH) assay for HBEGF detection in combination with a custom TUNEL IHC assay was performed using the automated Leica Bond™ platform (Leica Biosystems, Melbourne, Australia). The combination of ISH and IHC assays were optimized and performed at Advanced Cell Diagnostics (Newark, CA). To detect AAV-hDTR transduced cells a single-color probe for HBEGF was custom made and designed against the target region of 3 – 652 nt. The ISH was performed using the RNAscope®

2.5 LS Reagent Kit-RED (#322150) on 4 µm FFPE tissue sections according to the manufacturer's instructions followed by the TUNEL IHC as described above. RNA quality was evaluated for each sample with a probe specific to the housekeeping gene cyclophilin B (PPIB). A negative control background staining was evaluated using a probe specific to the bacterial dapB gene.

**Laser-capture microdissection**

FFPE lungs from AAV-stuffer (1 x $10^{11}$ vg) and AAV-hDTR (0.3 x $10^{11}$ vg) mice were sectioned at 10 µm thickness using a microtome, collected on polyethylene naphthalate membrane glass slides (UV-irradiated for 20 min for better adhesion of the tissue; Carl Zeiss Microscopy GmbH, Jena, Germany) and air dried over night at room temperature. Prior to LCM, sections on glass slides were deparaffinized using xylene (2x 3 min) and sequential ethanol washes (100%, 96%, 70%; 2x 1 min each) followed by H&E staining. Laser microdissection and pressure catapulting of ~3,000 cells was performed using a 20x objective under brightfield optics on a PALM MicroBeam system (Carl Zeiss Microscopy GmbH). The area equivalent to ~3,000 cells was estimated by counting and averaging the cell number of four areas in the region of interest. An amount of ~3,000 cells reflected an area of 220,000 – 290,000 $µm^2$ for the bronchial epithelium and 0.3 – 1.7 $mm^2$ for the alveolar epithelium (normal and infiltrated) sample depending. Isolated cells were collected in an adhesive cap (Carl Zeiss Microscopy GmbH).

**Protein extraction**

Microdissected tissue samples were processed as described recently (1). Briefly, samples were resuspended in 100 µL SDS extraction buffer (2% SDS in 300 mM Tris-HCl pH 8.0) and transferred into 0.5 mL Eppendorf Protein LoBind® tubes. Samples were boiled (25 min, 99 °C, 350 rpm) and sonicated for 20 cycles (QSonica Q700 with microplate horn, Newtown, CT; 30 s on, 30 s off) followed by heating for 2 h at 80 °C (500 rpm) and another sonication step (20 cycles). Reversibly oxidized cysteines were reduced with 10 mM DTT (30 min, 50 °C, 500 rpm) followed by alkylation of free thiols with 20 mM iodoacetamide (30 min, 22 °C, 500 rpm, in the dark).

**Protein digestion using the SP3 method**

Protein extracts were cleaned and digested with the SP3 method as described previously with modifications using the KingFisher Flex system (Thermo Scientific) (2, 3). Briefly, 20 µL of a 20 µg/µL SP3 bead stock (Sera-Mag SpeedBead carboxylate-modified magnetic particles; GE

Healthcare Life Sciences, Freiburg, Germany) and 300 µL acetonitrile (ACN; final concentration of 70%) were added to 100 µL of protein extract in the sample 96 deep-well plate. Proteins were bound to the beads (4.5 min medium speed, 30 s bottom mix) and washed two times each with 70% ethanol and ACN (500 µL; 30 s medium). Washing steps were performed without release of beads. After air-drying beads for 20 s beads with bound proteins were released into 100 µL 100 mM triethylammonium bicarbonate (TEAB; 1 min medium, 30 s bottom mix) followed by digestion for 8 h with 0.5 µg LysC/trypsin (37 °C, 1000 rpm). After centrifugation (10 min, 4000 g) beads were collected using the KingFisher Flex system and peptide samples were transferred into 0.5 mL tubes and dried *in vacuo* in a Concentrator plus (Eppendorf, Hamburg).

**TMT labelling**

Peptides were resuspended in 14 µL 100 mM TEAB. 100 µg TMTpro 16plex tags (Thermo Scientific) were dissolved in 6 µL anhydrous ACN and added to the corresponding samples followed by incubation for 2 h (22 °C, 600 rpm). Unreacted TMTpro tags were quenched with 5 µL 5% hydroxylamine (30 min). Labelled peptide samples belonging to the same batch were combined and dried *in vacuo*.

**High pH reversed phase fractionation**

TMTpro labelled peptides were fractionated using off-line high pH reversed phase chromatography. Dried samples were resuspended in 50 µL 5% formic acid, loaded onto a 2.1 x 150 mm XBridge BEH130 C18 column (3.5 µm, 130 Å; Waters) and separated on a Dionex Ultimate 3000 HPLC system with a flow rate of 0.17 mL/min. Solvents used were water (A), ACN (B) and 100 mM ammonium formate pH 9 (C). While solvent C was kept constant at 10%, solvent B started at 1%, increased to 10% in 3 min, 21.5% in 2 min, 50% in 13 min and 90% in 1 min, was kept at 90% for further 4 min followed by returning to starting conditions and re-equilibration for 7 min. Peptides were collected every 52 s into total 24 fractions, which were concatenated into 8 fractions and subsequently dried *in vacuo*. Peptides were redissolved in 3% ACN/0.1% TFA and analysed by LC-MS.

**LC-MS analysis**

TMTpro labelled fractions were separated on a Dionex RSLCnano HPLC and analysed on an Orbitrap Eclipse Tribrid mass spectrometer in combination with the FAIMS Pro Interface (Thermo

Scientific). Peptides were loaded onto a 100 µm × 2 cm Acclaim PepMap-C18 trap column (5 µm, 100 Å) for 6 min with 3% ACN/0.1% TFA and a constant flow of 10 µL/min and separated on a 75 µm × 75 cm EASY-Spray C18 column (2 µm, 100 Å; Thermo Scientific) at 50 °C with a flow rate of 300 nL/min. Solvents used were 0.1% formic acid (A) and 80% ACN/0.1% formic acid (B). Samples were separated over a multi-step gradient from 10% to 20% B in 24 min, to 28% in 37 min and 40% in 52 min with a total measurement time of 150 min. The spray was initiated by applying 1.7 kV to the EASY-Spray emitter. The ion transfer capillary temperature was set to 275 °C and the radio frequency of the ion funnel to 30%. Data were acquired under the control of the Xcalibur software in a data-dependent mode using advanced peak detection. The FAIMS Pro interface was used in standard resolution mode without gas flow. Two compensation voltages with asymmetric cycle times were applied (-40/-60 V for 1.2/0.8 s). The full scan was acquired in the orbitrap covering the mass range of *m/z* 375-1,500 with a mass resolution of 120,000, a normalized automatic gain control (AGC) target of 100% and a maximum injection time of 50 ms. Only precursor ions with charges between 2 and 7, a minimum intensity of $1x10^4$ and a minimum of 70% of signal within an isolation window of *m/z* 0.7 were selected for fragmentation using collision-induced dissociation in the ion trap with 34% collision energy. MS2 spectra were acquired with a rapid ion trap scan rate, a normalized AGC target of 100% and a maximum injection time of 105 ms using a dynamic exclusion of 60 s. The real-time search algorithm was applied in SPS mode in the cycle with FAIMS CV -40 using the following settings: Uniprot mouse database, carbamidomethylation (Cys) and TMTpro (N-term/Lys) as fixed modifications, oxidation (Met) as variable modification, trypsin (full), one missed cleavage and one variable modification per peptide and a maximum search time of 35 ms. An Xcorr of 1.4 and a mass tolerance of 15 ppm for precursor ions were used as scoring thresholds. For the MS3 analysis 10 (CV -40) or 7 (CV -60) fragment ions were co-isolated using synchronous precursor selection in a window of *m/z* 2 and further fragmented with an HCD collision energy of 45%. The resulting fragments were analysed in the orbitrap with a resolution of 50,000, a normalized AGC target of 200% and a maximum injection time of 150 ms.

**Data analysis**

Acquired tandem mass spectra were searched against the UniProtKB/Swiss-Prot mouse database (downloaded in December 2019, containing 17,025 protein sequences) using the Sequest search engine in Proteome Discoverer 2.4 (Thermo Scientific). Trypsin was specified as the cleavage enzyme, allowing up to two missed cleavages and a mass tolerance of 10 ppm for precursor ions and 0.6 Da for fragment ions. Carbamidomethylation of cysteine and TMTpro-modification of the N-terminus and lysine were used as fixed modifications and oxidation of methionine and acetylation of the protein N-terminus as variable modifications. Using the percolator node, a decoy-based false discovery rate (FDR) of 1% was applied to peptide-spectrum matches. Additionally, an FDR of 1% was applied to peptide and protein identifications (high confidence). For the quantification of TMTpro reporter ion signals from unique peptides correction of isotopic impurities, an average reporter signal-to-noise ratio of 2, an SPS mass matches threshold of 65% and a co-isolation threshold of 100% were applied. Contaminants were excluded and only master proteins, proteins with high confidence and proteins with minimum 2 unique peptides per protein were considered for further analysis. TMTpro batches were analysed in independent database searches. All protein quantities (summed TMTpro reporter signal-to-noise ratios of corresponding peptides) of samples within a batch were median-normalized to address differences in sample loadings. Additionally, both alveoli batches (normal tissue hDTR vs stuffer and hDTR infiltrated vs normal) were searched together followed by median normalization and another normalization step to address the variation between batches. The median of protein quantities of the seven hDTR normal tissue samples was calculated for each batch and used to calculate a normalization factor as described in Griesser et al. (1). Principal component analysis and volcano plots were generated with Perseus (version 1.6.7.0) (4) and hierarchical clusters were created using Instant Clue (version 0.5.3) (5). Functional enrichment and pathway analysis was performed with the DAVID tool (6, 7).

#### Supplemental tables

**Table S1:** Mean cytokine concentrations in pg/ml detected in the BALF of AAV-stuffer (1 x $10^{11}$ vg) and AAV-hDTR treated animals 24 h after intratracheal administration of 100 ng DT (n=5-7). Cytokines were measured with the 35-plex MSD panel. SD – standard deviation.

| Cytokine | Stuffer control | | 0.03 x $10^{11}$ vg | | 0.1 x $10^{11}$ vg | | 0.3 x $10^{11}$ vg | | 1 x $10^{11}$ vg | |
|---|---|---|---|---|---|---|---|---|---|---|
| pg/ml | Mean | SD | Mean | SD | Mean | SD | Mean | SD | Mean | SD |
| EPO | 11.92 | 8.47 | 1.24 | 2.13 | 3.64 | 8.91 | 5.18 | 7.40 | 1.60 | 3.93 |
| GM-CSF | 56.78 | 47.96 | 0.83 | 0.76 | 36.64 | 86.44 | 0.34 | 0.34 | 0.12 | 0.18 |
| IFNg | 0.004 | 0.012 | 0.002 | 0.006 | 0.065 | 0.090 | 0.671 | 0.480 | 0.484 | 0.355 |
| IL12p70 | 0.000 | 0.000 | 0.000 | 0.000 | 0.000 | 0.000 | 0.000 | 0.000 | 0.000 | 0.000 |
| IL1b | 0.007 | 0.018 | 0.079 | 0.131 | 0.037 | 0.058 | 0.211 | 0.173 | 0.036 | 0.087 |
| IL2 | 0.000 | 0.000 | 0.000 | 0.000 | 0.000 | 0.000 | 0.000 | 0.000 | 0.000 | 0.000 |
| IL5 | 0.000 | 0.000 | 6.695 | 7.146 | 4.352 | 5.737 | 1.912 | 1.473 | 0.346 | 0.537 |
| IL6 | 355.0 | 214.8 | 1921.1 | 1657.2 | 812.8 | 851.1 | 598.3 | 399.0 | 242.8 | 166.0 |
| KC/GRO | 197.6 | 84.7 | 28.8 | 27.2 | 74.8 | 139.5 | 14.3 | 7.1 | 7.2 | 2.9 |
| TNFa | 0.000 | 0.000 | 0.437 | 0.641 | 4.051 | 7.119 | 18.354 | 8.970 | 16.580 | 7.440 |
| IL-10 | 0.000 | 0.000 | 0.002 | 0.004 | 0.095 | 0.147 | 0.231 | 0.422 | 0.159 | 0.255 |
| IL-13 | 0.108 | 0.285 | 0.562 | 1.147 | 0.000 | 0.000 | 0.072 | 0.191 | 0.548 | 0.891 |
| IL-15 | 22.0 | 28.3 | 19.0 | 20.8 | 1.2 | 3.0 | 1.5 | 2.3 | 17.9 | 9.1 |
| IL-17F | 5.4 | 8.1 | 7.9 | 5.7 | 0.0 | 0.0 | 0.0 | 0.0 | 8.1 | 6.4 |
| IL-23 | 0.049 | 0.129 | 0.000 | 0.000 | 0.001 | 0.003 | 0.009 | 0.021 | 0.053 | 0.119 |
| IL-31 | 0.000 | 0.000 | 0.000 | 0.000 | 0.328 | 0.804 | 1.481 | 3.919 | 0.000 | 0.000 |
| IL-33 | 0.868 | 0.569 | 1.932 | 1.420 | 1.924 | 0.793 | 2.219 | 0.869 | 2.457 | 0.985 |
| IL-4 | 0.013 | 0.025 | 0.005 | 0.006 | 0.001 | 0.002 | 0.019 | 0.034 | 0.030 | 0.032 |
| VEGF | 2.945 | 1.328 | 6.963 | 4.207 | 9.872 | 5.044 | 7.042 | 5.114 | 2.396 | 0.772 |
| IL-21 | 1.555 | 2.504 | 9.492 | 19.433 | 3.031 | 7.060 | 4.781 | 4.322 | 19.382 | 11.966 |
| IL-22 | 0.285 | 0.276 | 0.568 | 1.074 | 0.260 | 0.309 | 1.002 | 0.614 | 1.346 | 0.830 |
| IL-17C | 0.427 | 0.737 | 0.000 | 0.000 | 0.000 | 0.000 | 1.012 | 1.547 | 0.959 | 1.137 |
| IL-12 | 62.3 | 25.1 | 351.1 | 273.1 | 921.9 | 542.2 | 3095.0 | 1143.5 | 3227.9 | 1023.6 |
| IL-17A | 0.058 | 0.074 | 0.073 | 0.120 | 0.032 | 0.044 | 0.301 | 0.163 | 0.310 | 0.162 |
| IL-16 | 198.7 | 69.9 | 170.3 | 125.9 | 227.0 | 83.1 | 340.8 | 104.7 | 355.5 | 67.6 |
| IL-17E/ IL-25 | 0.165 | 0.303 | 0.421 | 0.875 | 0.000 | 0.000 | 0.038 | 0.100 | 0.247 | 0.553 |
| IL-17 A/F | 0.000 | 0.000 | 0.000 | 0.000 | 2.249 | 3.081 | 1.351 | 1.689 | 0.256 | 0.477 |
| IP-10 | 13.8 | 7.6 | 410.6 | 377.7 | 1045.6 | 563.8 | 2888.9 | 382.9 | 2399.8 | 662.4 |
| MCP-1 | 7.9 | 2.9 | 58.6 | 51.5 | 60.5 | 43.4 | 160.3 | 35.6 | 90.1 | 39.3 |
| MIP-1a | 5.8 | 2.4 | 6.6 | 5.4 | 8.4 | 3.5 | 15.0 | 3.1 | 10.8 | 2.5 |
| MIP-1b | 26.4 | 11.8 | 49.7 | 41.6 | 98.9 | 48.8 | 184.8 | 31.7 | 131.3 | 34.0 |
| MIP-2 | 131.3 | 67.0 | 21.4 | 13.5 | 51.1 | 62.2 | 47.6 | 16.2 | 37.9 | 9.6 |
| MIP-3a | 54.1 | 26.3 | 66.1 | 47.4 | 93.5 | 46.1 | 237.2 | 106.9 | 213.9 | 45.7 |

#### Supplemental figures

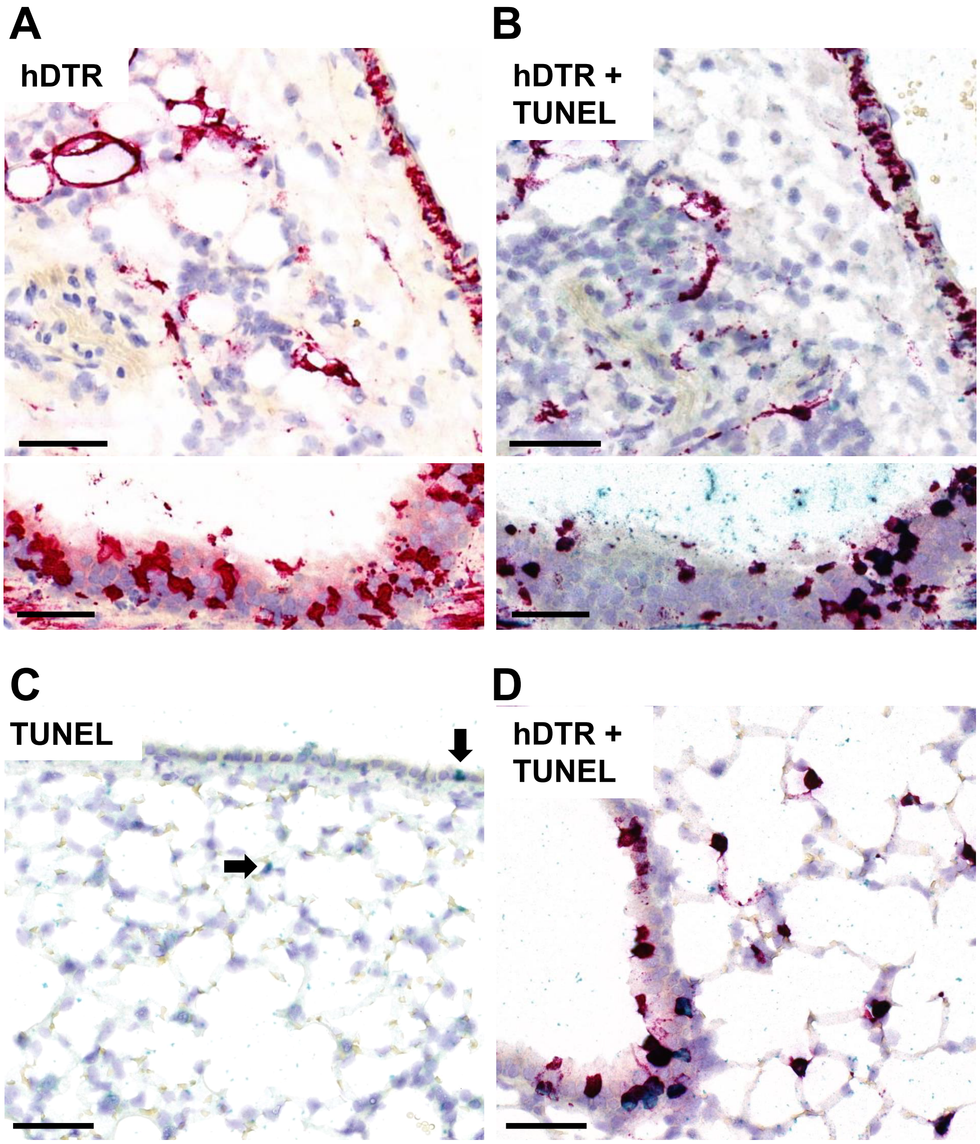


**Figure S1**: Representative images of co-staining for hDTR mRNA expression (*in situ* hybridization - ISH, red) and apoptosis (TUNEL, green) 24 h after intratracheal administration of 100 ng DT. (A) Single ISH showed hDTR mRNA expression (red) detected in bronchial and alveolar epithelial cells in AAV-hDTR animals (1 x $10^{11}$ vg). (B) On a subsequential section ISH for hDTR mRNA expression (red) was performed followed by TUNEL-staining (green) leading to purple staining in colocalized areas. (C) Bronchial epithelium and parenchyma of an AAV-stuffer treated animal (1 x $10^{11}$ vg) showed only occasional TUNEL positive cells (arrows). (D) Colocalization (purple) of hDTR mRNA expression and apoptosis in bronchial and alveolar epithelial cells of an AAV-hDTR treated animal (1 x $10^{11}$ vg). Scale bars, 50 µm.

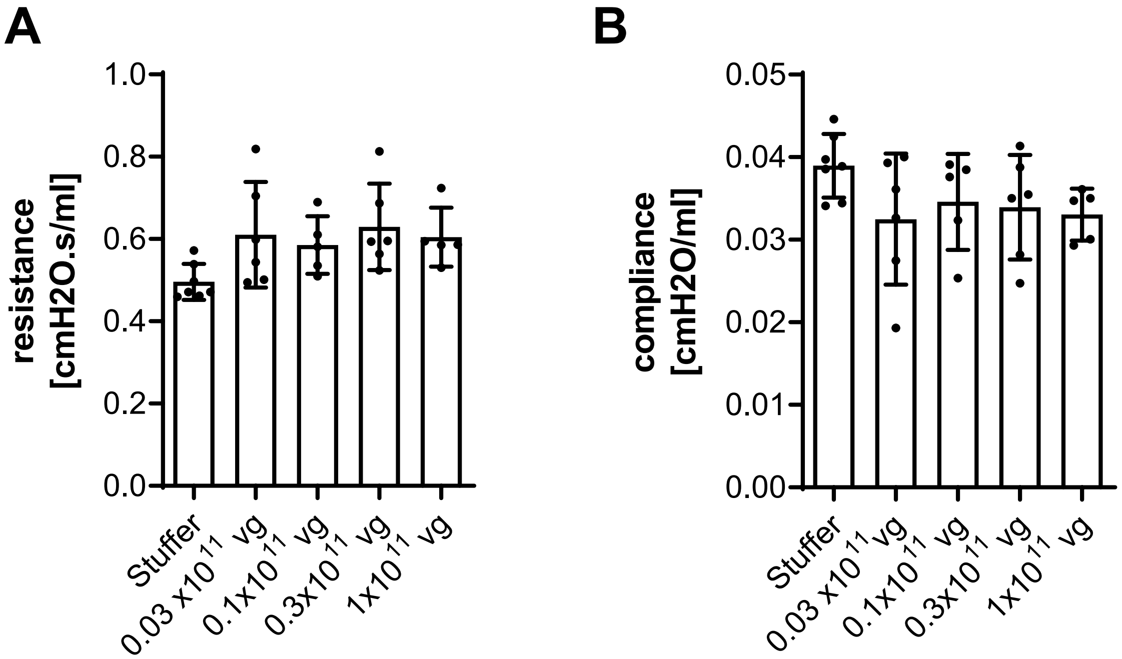


**Figure S2**: Representative data of lung function measurements in AAV-stuffer (1 x $10^{11}$ vg) and AAV-hDTR treated animals 24 h after administration of the highest dose of DT (200 ng). (A) Resistance of the respiratory system as well as (B) compliance were measured using the flexiVent system (n=5-7).

A B

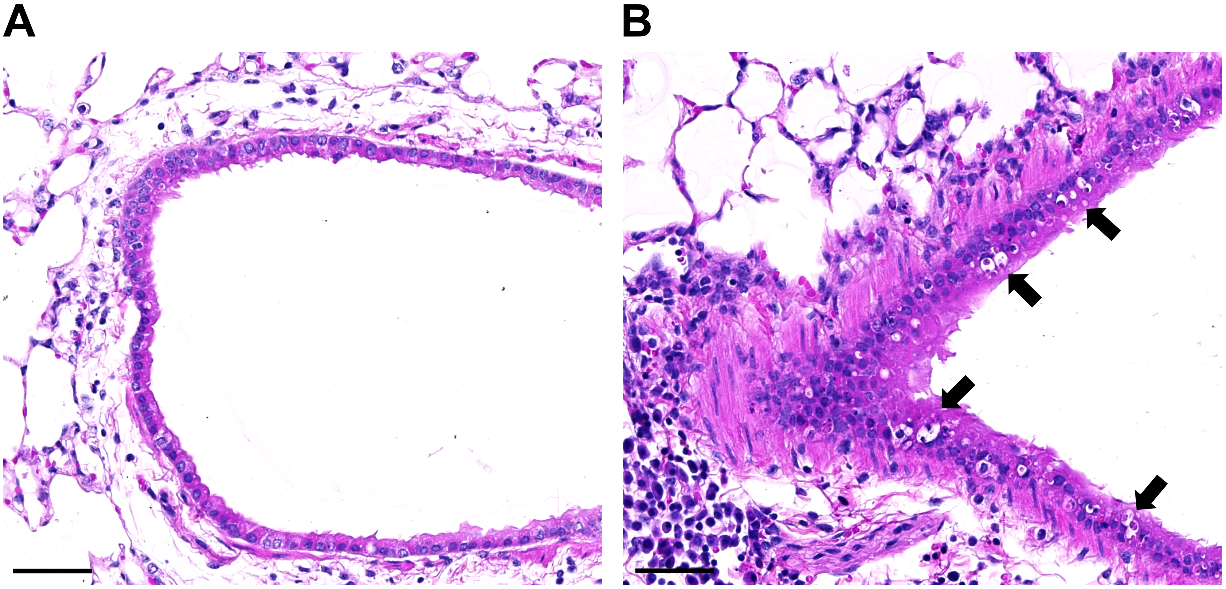

**Figure S3**: Representative images of H&E-stained lung sections 24 h after administration of 100 ng DT. (A) Normal bronchial epithelium of an AAV-stuffer treated animal (1 x $10^{11}$ vg). (B) Pyknotic nuclei and apoptotic bodies (arrows) in bronchial epithelial cells of an AAV-hDTR treated animal (1 x $10^{11}$ vg). Also, inflammatory influx surrounding the bronchus can be seen. Scale bars, 50µm.

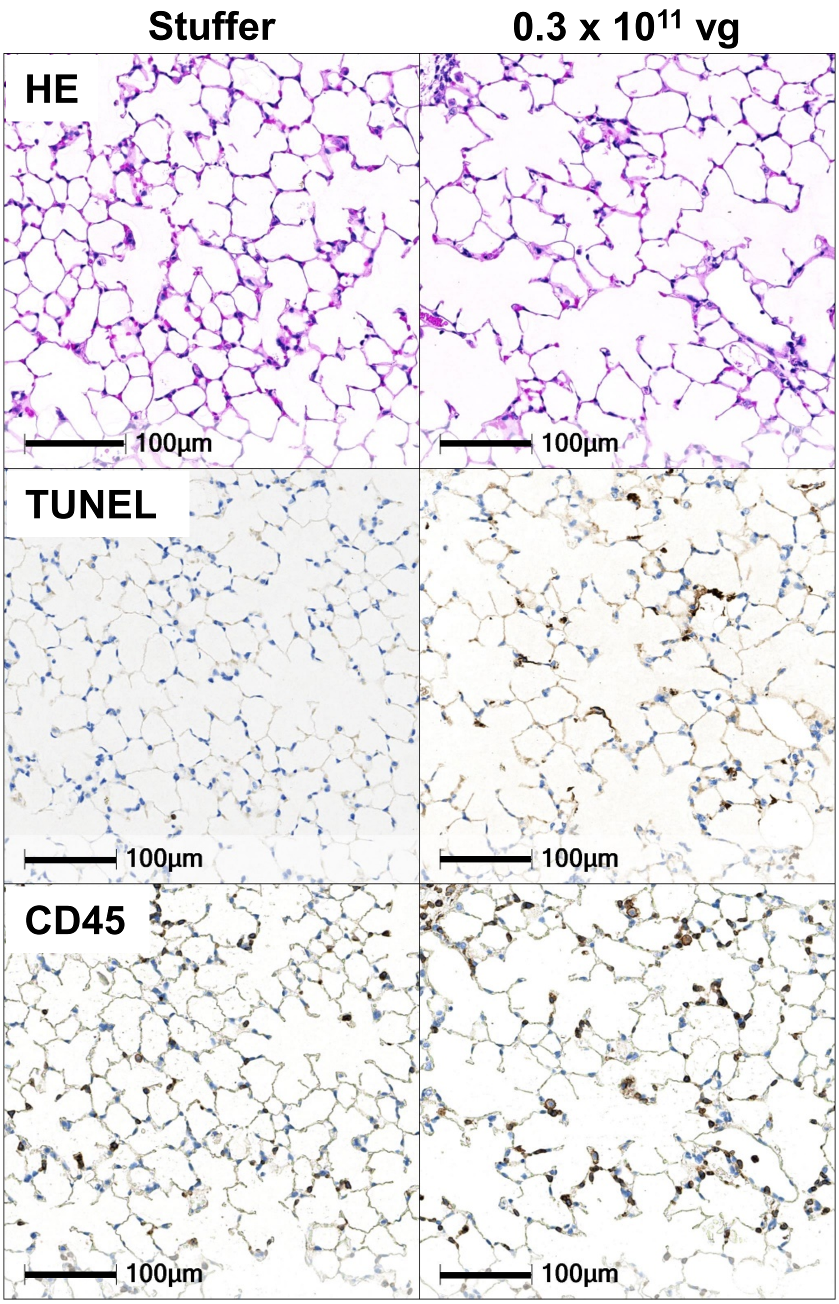


**Figure S4**: Representative images of H&E-, TUNEL- and CD45-stained sequential lung sections showing apoptotic alveolar epithelial cells (middle) and resulting leukocyte influx (bottom) in AAV-hDTR treated animals (0.3 x $10^{11}$ vg) compared to AAV-stuffer treated animals (1 x $10^{11}$ vg) 24 h after intratracheal administration of 100 ng DT. Alveolar airways contain only occasional inflammatory cells as bronchoalveolar lavage and agarose instillation of lungs were performed before further tissue processing.

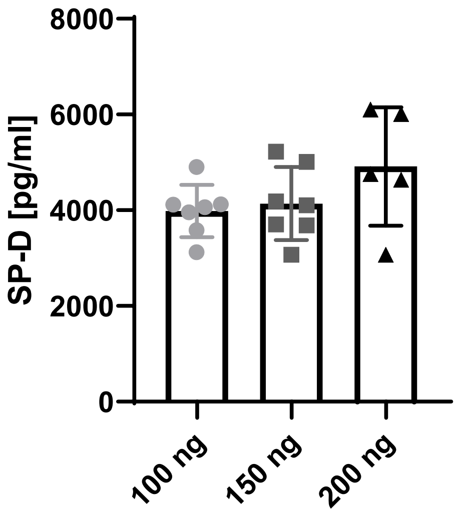


**Figure S5**: SP-D levels detected in plasma of AAV-stuffer treated animals (1 x $10^{11}$ vg) 24 h after intratracheal administration of indicated diphtheria toxin doses (n=5-7).

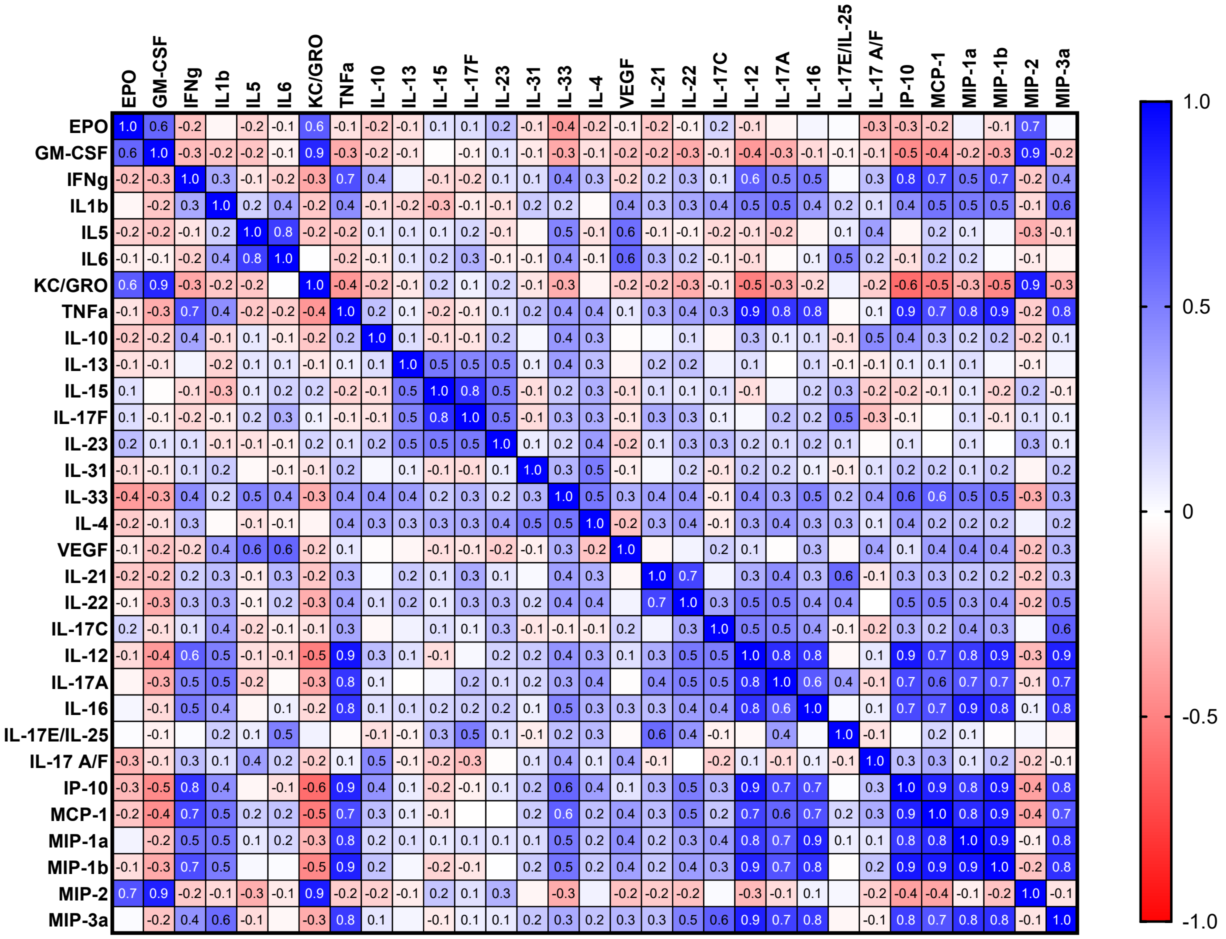


**Figure S6**: Pearson correlation matrix of detected cytokines as stated in Table S1.

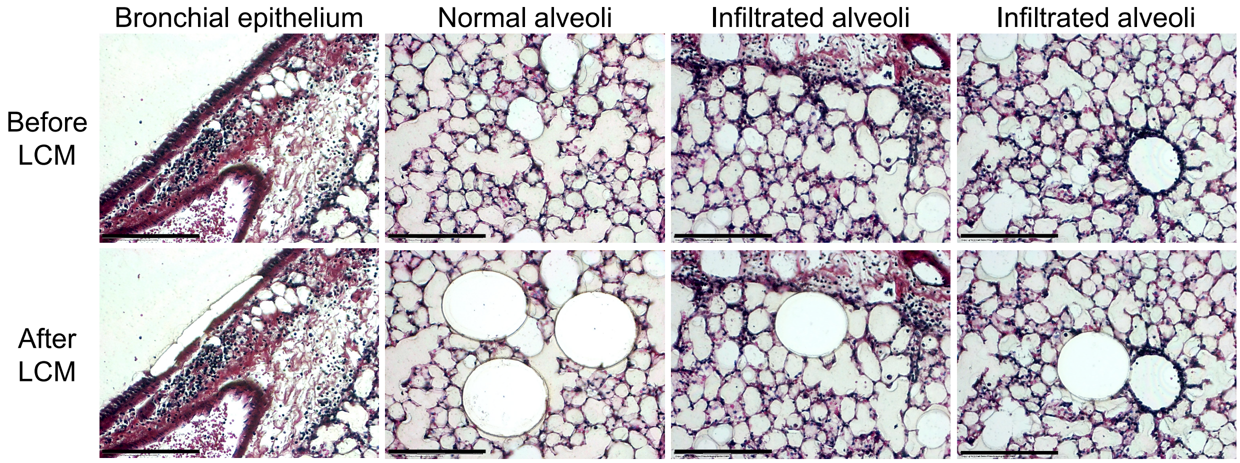


**Figure S7**: Laser-capture microdissection (LCM) of different lung regions from formalin-fixed and paraffin-embedded tissue. Representative images show the region of interest before and after LCM. Images were acquired from H&E-stained lung tissue sections (10 µm thickness) with a 20x objective using the Zeiss PALM MicroBeam system. Scale bars, 150 µm.

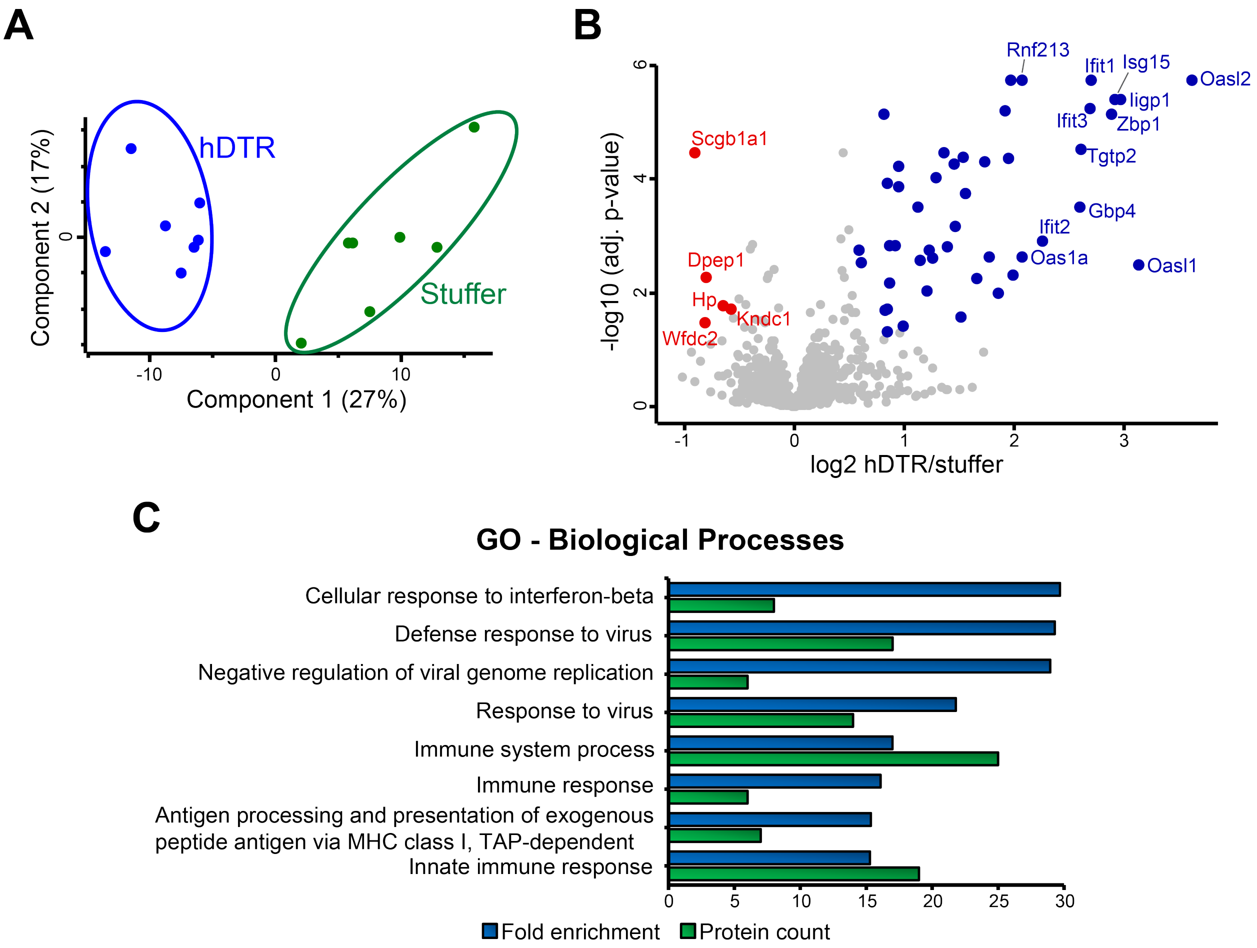


**Figure S8**: Proteomic changes in bronchial epithelium upon acute epithelial injury. (A) Principal component analysis scores plot of log2-transformed protein quantities of AAV-hDTR (0.3 x $10^{11}$ vg) and AAV-stuffer (1 x $10^{11}$ vg) mice 24 h after intratracheal instillation with 100 ng DT. (B) Volcano plot representing the negative log10-transformed Benjamini Hochberg (BH) adjusted p-values vs. the log2 fold changes in protein quantities of AAV-hDTR mice compared to AAV-stuffer mice (n=7; two-tailed, equal variance Student's t-test). Proteins with fold changes ≥1.5 (blue) or ≤1.5 (red) are displayed in colour. (C) Overrepresented Gene Ontology (GO) Biological Processes annotated to proteins significantly upregulated (fold change ≥1.5, BH-adjusted p-value <0.05) in AAV-hDTR compared to AAV-stuffer mice. Bronchial epithelium-specific proteome was used as background (Fisher's exact test, BH-corrected p-value <0.001, protein count >5). Protein quantities are the summed TMTpro reporter ion signal-to-noise ratios of corresponding peptides.
